# Macrophage-derived reactive oxygen species promote *Salmonella* aggresome formation contributing to bacterial antibiotic persistence

**DOI:** 10.1101/2025.06.03.657582

**Authors:** Xiao Chen, Kefan Fang, Bo Li, Yingxing Li, Yuehua Ke, Weixin Ke, Tian Tian, Yifan Zhao, Linqi Wang, Jing Geng, Mark C. Leake, Fan Bai

## Abstract

In this study, we reveal that macrophage-derived reactive oxygen species (ROS) can trigger the rapid formation of *Salmonella* aggresomes, which substantially contribute to the increased frequency of persisters induced by phagocytosis. *Salmonella* containing aggresomes exhibited a dormant phenotype characterized by reduced adenosine triphosphate (ATP) levels and decreased metabolic activity. Furthermore, these dormant bacteria showed upregulated expression of *Salmonella* pathogenicity island 1 (SPI-1)-encoded type III secretion system (T3SS)-related genes, followed by later expression of SPI-2 T3SS-related genes when macrophages ROS production declined. Our results demonstrate that *Salmonella* containing aggresomes can enter a dormant state to escape antibiotic attack, while crucially maintaining the ability to resuscitate when the stress environment is improved. Research on bacterial aggresomes could potentially provide therapeutic strategies to combat bacterial antibiotic persistence.

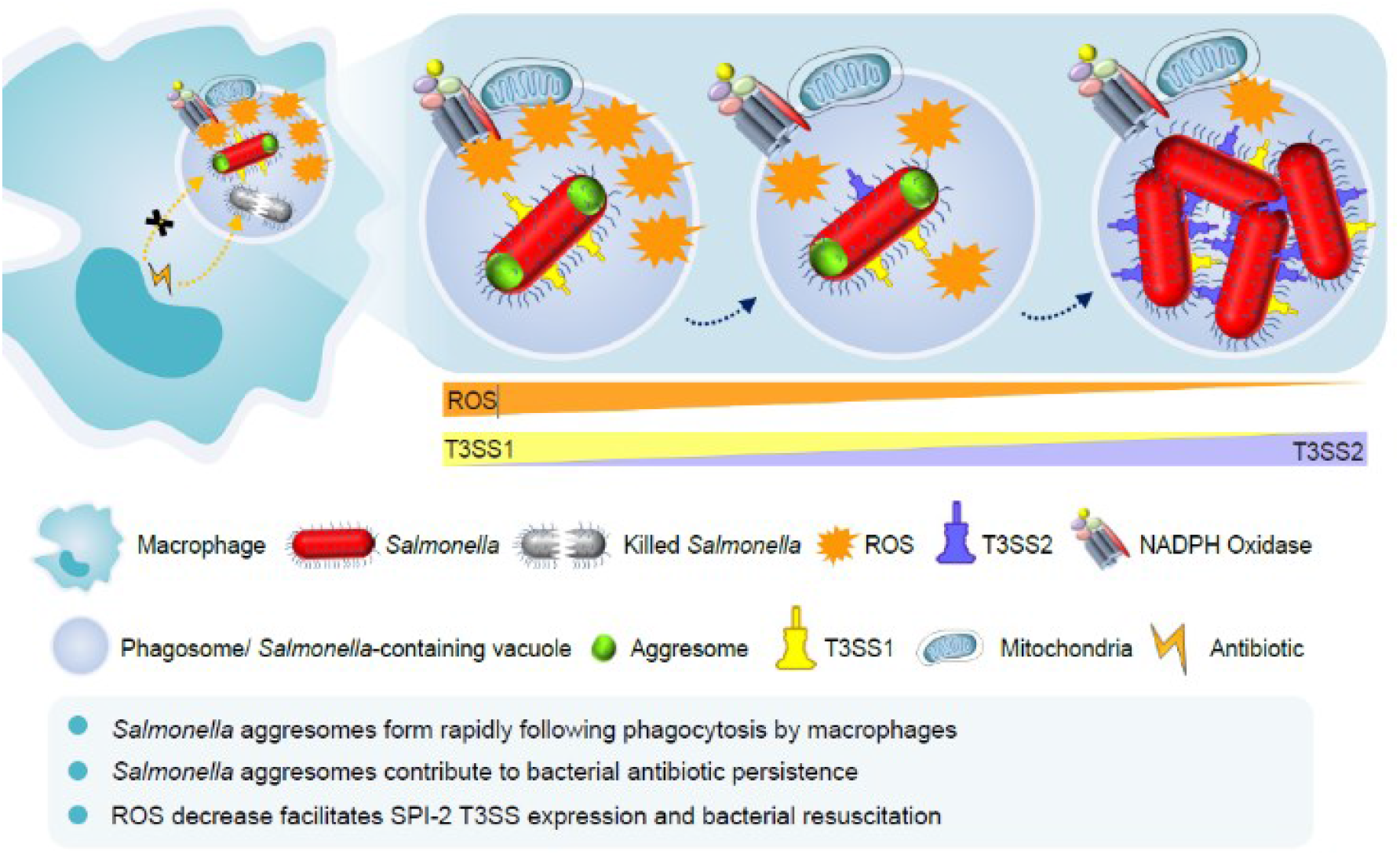

---

Bacterial persisters represent a subpopulation of phenotypic variants that survives antibiotic attack and subsequently resuscitate [1], serving as reservoirs for recurrent infections [2]. Bacterial dormancy is regarded as a prominent theory in elucidating persister formation, as dormant bacteria show decreased metabolic activity and suppressed proliferation rates [2, 3]. Several molecular mechanisms are involved in bacterial dormancy [4]. For example, reactive oxygen species (ROS) deactivate the tricarboxylic acid (TCA) cycle, resulting in a reduction in respiration and adenosine triphosphate (ATP) generation [5]. In addition, the activation of type I toxin TisB and HokB cause ATP leakage, leading to cell dormancy [6]. The type II toxin HipA triggers cell dormancy by phosphorylating the essential translation factor EF-Tu [7].

Upon host entry, pathogenic bacteria encounter multiple stresses including acidic pH, nutrient limitation, ROS and reactive nitrogen species (RNS) [8]. A previous study reported that following macrophage phagocytosis, the proportion of *Salmonella* persisters can greatly increase depending on the expression of several type II toxin-antitoxin (TA) genes induced by vacuolar acidification and (p)ppGpp synthesis [9]. However, several independent follow-up studies have argued that deletion of 10 type II TA modules does not affect persister cell levels [10, 11]. Moreover, type II toxins are not always activated under diverse stress conditions [11]. Therefore, the critical mechanism mediating macrophage-induced antibiotic persistence remains unclear.

Recently, several studies have elucidated the correlation between bacterial aggresomes and dormancy [3, 12, 13]. Bacterial aggresomes contain many proteins which are vital for cellular functions associated with carbon metabolism, oxidative phosphorylation, transcription, and translation [3]. The formation of bacterial aggresomes is driven by liquid–liquid phase separation (LLPS) [12]. Here, we explore the relationship between aggresomes formation and macrophage-induced bacterial antibiotic persistence in *Salmonella*.

## RESULTS AND DISCUSSION

### *Salmonella* aggresomes form rapidly following phagocytosis by macrophages

*Salmonella Typhimurium* (*S. Typhimurium*) SL1344 HslU was genetically fused with EGFP to enable visualization of bacterial protein aggresomes (Figure 1A, S1A,B). Following phagocytosis of fluorescently labeled *Salmonella* at a multiplicity of infection (MOI) of 100 by RAW264.7 macrophages, distinct HslU-EGFP foci were detected *in situ* as early as 0.5 hours post infection (h.p.i.) (Figure 1B). The percentage of bacterial cells containing HslU-EGFP foci was 19.7% (Figure 1C). Moreover, bacteria showing HslU-EGFP foci were also observed in human macrophages differentiated from monocytic leukemia cells (THP-1) and immortalized murine bone marrow-derived macrophages (iBMDMs) (Figure 1C, S1C). Both exponential-phase and stationary-phase bacteria formed protein aggresomes upon macrophage infection, confirming that the formation of intracellular bacterial aggreomes is independent of bacterial growth state. In addition, following macrophage phagocytosis, aggresomes labelled by FITC staining are formed in other bacteria species such as *Shigella flexneri* (*S. flexneri*) and *Mycobacterium smegmatis* (*M. smegmatis*) (Figure S1D). Following the invasion of macrophages, *Salmonella* reside in phagosomes, which undergo a series of maturation steps [8]. To further determine the stage of phagosome maturation at which these *Salmonella* form aggresomes, immunofluorescence staining of early endosomes and lysosomes was performed. Phagosomes containing *Salmonella* with aggresomes colocalized with both early endosomes (EEA1) and lysosomes (LAMP1) (Figure S1E), suggesting that gradually matured phagosomes featured by a hostile environment promote the formation of *Salmonella* aggresomes.

**FIGURE 1.**
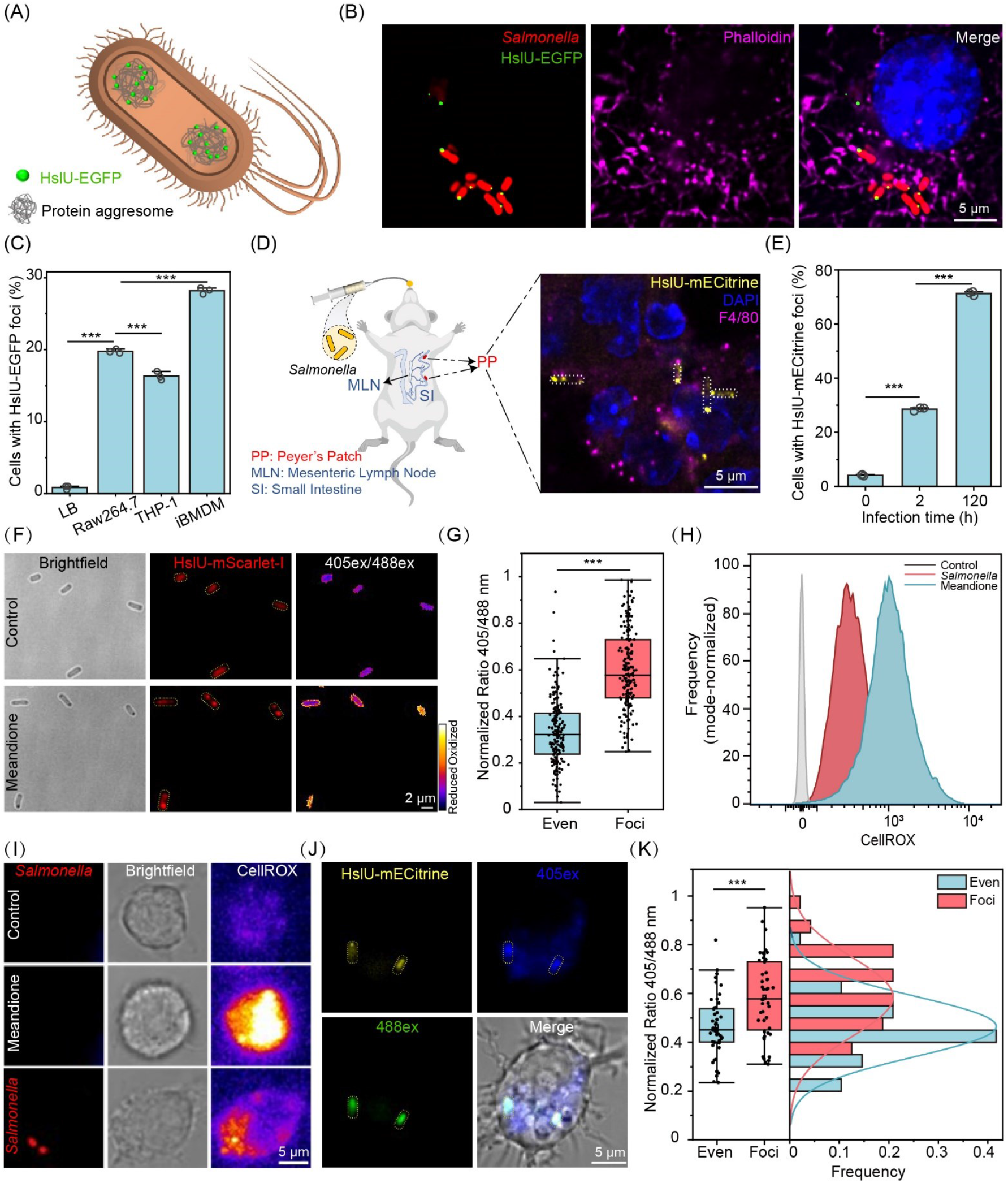
The presence of *Salmonella* aggresomes within macrophages. (A) Schematic of fluorescently labeled bacteria. (B) Immunofluorescence microscopy imaging of bacterial aggresomes within macrophages. The actin cytoskeleton was stained with phalloidin (magenta), and nuclei were stained with DAPI (blue) (scale bar, 5 μm). (C) Percentage of *Salmonella* in the stationary phase possessing HslU-EGFP foci after internalization by RAW264.7 macrophages, THP-1 cells, and iBMDMs. Each data point represented an independent biological replicate (*n* = 3); 200 bacteria were analyzed per replicate. (D) Representative confocal micrograph of *Salmonella* aggresomes in infected murine PPs at 120 h.p.i. The cryosection was labeled with an antibody to the macrophage marker F4/80 (magenta). Aggresomes were labeled by HslU-mECitrine (yellow) (scale bar, 5 μm). (E) Percentage of *Salmonella* cells with HslU-mECitrine foci in murine PPs at 2 and 120 h.p.i.; *n* = 3. (F) Images of *Salmonella* showing aggresomes and ROS levels following challenge with 160 μM menadione. Bacterial aggresomes were labeled with HslU-mScarlet-I, and the 405ex/488ex ratio was used as a measurement of roGFP2_oxidized_/roGFP2_reduced_ (scale bar, 2 μm). (G) Average ROS levels of bacteria with and without aggresomes. (H, I) *Salmonella* infection promoted ROS production in macrophages. (H) Representative histograms of CellROX. (I) Fluorescence microscopy images of ROS production in macrophages. *Salmonella* were labeled with mCherry (scale bar, 5 μm). (J) Live-cell imaging of bacterial ROS levels and aggresome formation in *Salmonella* within RAW264.7 cells at 1.5 h.p.i. (scale bar, 5 μm). (K) Histogram of the ROS levels within intracellular bacteria with and without aggresomes. C and E were assessed using one-way ANOVA followed by Bonferroni post hoc test; G and K were assessed using two-tailed unpaired *t*-test. Error bars indicate standard deviation; * *p* < 0.05, ** *p* < 0.01, and *** *p* < 0.001.

Next, C57BL/6 mice were intragastrically inoculated with *Salmonella* whose HslU protein was labeled by acid-tolerant mECitrine [14]. Immunofluorescence imaging of F4/80 was performed on cryosections of Peyer’s patches (PPs) from *Salmonella*-infected mice, and *Salmonella* with HslU-mECitrine foci were observed within murine macrophages (Figure 1D). Subsequently, the PPs were lysed, and the released *Salmonella* cells were subjected to microscopy imaging. As early as 2 h.p.i., 29% of *Salmonella* cells in the PPs displayed HslU-mECitrine foci, and at 120 h.p.i., the proportion of cells with fluorescence foci increased dramatically to 71% (Figure 1E). Whether bacterial aggresomes can form in patients with clinical infectious disease remains unclear and requires further investigation.

### Macrophage-derived ROS induces the formation of *Salmonella* aggresomes

To elucidate the driving force for the formation of bacterial aggresomes in macrophages, we evaluated the *ex vivo* effect of different phagosomal stress factors [8]. The results revealed that ROS substantially promoted the formation of *Salmonella* aggresomes (Figure 1F), whereas amino acid starvation induced by serine hydroxamate (SHX) and an acidic environment had no effect (Figure S1F,G). Then the ROS stress levels encountered by *Salmonella* were quantitated using the redox-sensitive biosensor roGFP2 [15]. Following exposure of *Salmonella* to 160 μM menadione, ROS levels in bacterial cells with fluorescently labeled HslU foci were 1.8-fold higher than those in cells lacking fluorescent foci (Figure 1F,G), suggesting a positive correlation between ROS levels and aggresome formation.

Next, we investigated the relationship between ROS levels and *Salmonella* aggresome formation within macrophages. An increase in CellROX intensity was observed by both flow cytometry and fluorescence imaging after infection, indicating that macrophages generate higher levels of ROS in response to *Salmonella* invasion (Figure 1H,I). We also demonstrated a correlation between the proportion of *Salmonella* exhibiting protein aggresomes (Figure 1C, S1C) and ROS level variations in different macrophage types (Figure S1H,I). Subsequently, we quantified the ROS stress levels encountered by *Salmonella* inside macrophages by using roGFP2 and found that *Salmonella* possessing aggresomes were subjected to a 1.3-fold higher levels of oxidative stress (Figure 1J,K). Moreover, treatment of macrophages with ROS inhibitors reduced the number of aggresome-positive *Salmonella* (Figure S1J,K). Additionally, C57BL/6 wild-type and ROS-deficient mice (*Cybb*^*-/-*^) mice were intragastrically inoculated with *Salmonella* to initiate acute systemic infection. Flow cytometry revealed reduced ROS levels in *Cybb*^-/-^ splenic macrophages compared to wild-type controls (Figure S2G,H). Concurrently, the proportion of *Salmonella* containing aggresomes in *Cybb*^-/-^ mice was lower in spleen, liver, mesenteric lymph nodes (MLNs), and PPs at 48 h.p.i. (Figure S1L).

### *Salmonella* aggresomes contribute to macrophage-induced bacterial antibiotic persistence

The persister ratio of *Salmonella* internalized by macrophages exhibited a 10-fold increase compared to uninfected LB-cultured controls (Figure 2A–C), suggesting bacterial aggresomes may enhance antibiotic persistence. To further verify this point, the bacterial antibiotic killing and resuscitation process was monitored by time-lapse fluorescence imaging. The result showed that ampicillin (100 μg/mL, 10 × MIC) rapidly lysed *Salmonella* without protein aggresomes, whereas *Salmonella* harboring HslU-EGFP foci entered a non-proliferative state and resumed growth in fresh LB medium (Figures 2D,E). Statistical analysis showed that 67.9% of persisters were derived from bacteria with aggresomes (Figure 2F). Considering that only 19.7% of phagocytosed *Salmonella* contained aggresomes (Figure 1C), the probability of persisters originating from cells possessing aggresomes was 8.6-fold higher than bacteria without aggresomes (Figure 2G).

**FIGURE 2.**
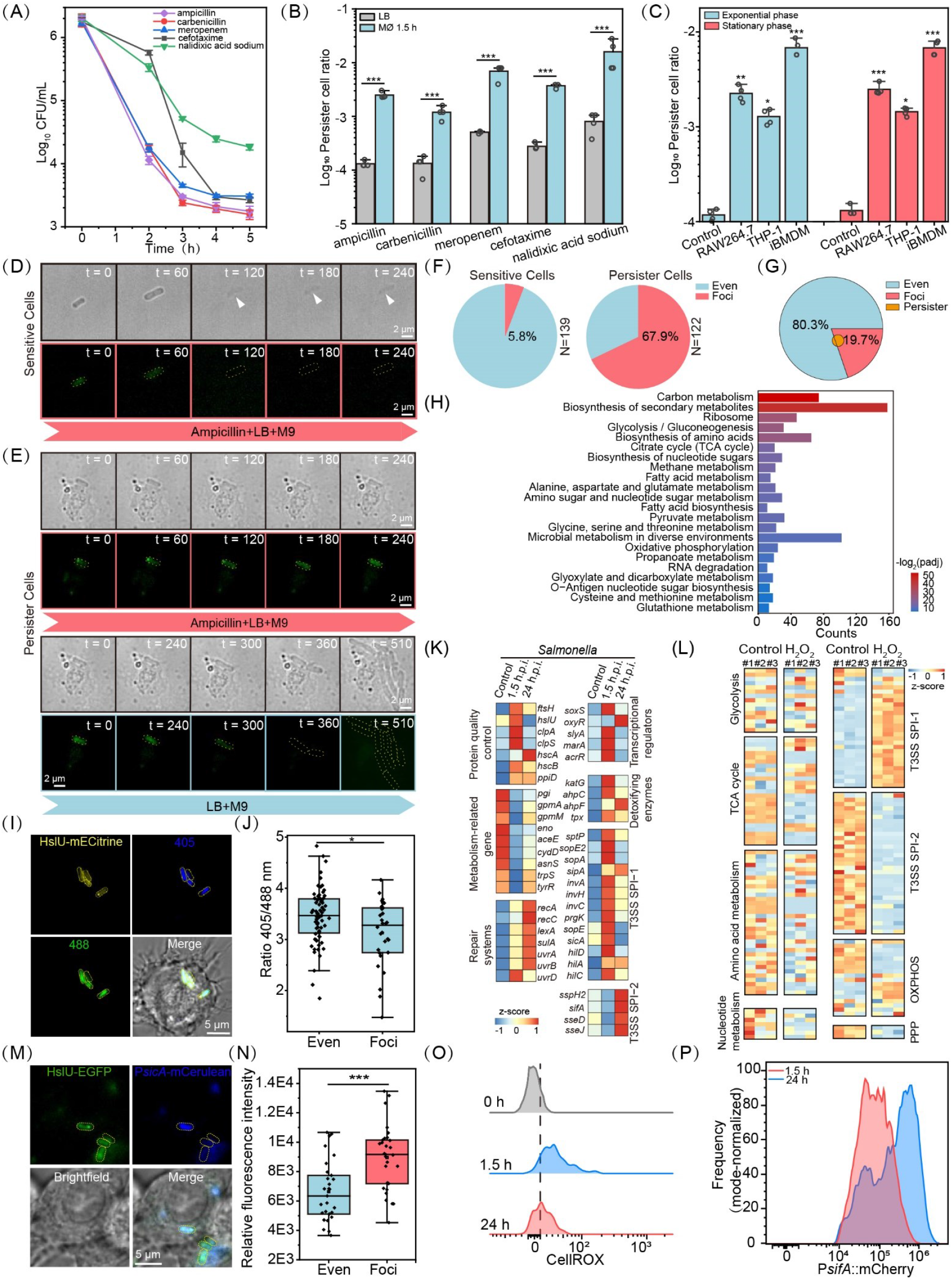
Protein aggresomes within *Salmonella* contribute to macrophage-induced bacterial antibiotic persistence. (A) Time-kill curves for *Salmonella* released from RAW264.7 macrophages exposed to 100 μg/mL ampicillin, 100 μg/mL carbenicillin, 5 μg/mL meropenem, 10 μg/mL cefotaxime, and 100 μg/mL nalidixic acid sodium for the indicated lengths of time. (B) Log percentage survival of LB medium-grown bacteria or bacteria at 1.5 h following phagocytosis by RAW264.7 macrophages after 5 h of treatment with ampicillin, carbenicillin, meropenem, cefotaxime, and nalidixic acid sodium treatment; *n* = 4. (C) Log percentage survival of LB medium-grown bacteria or bacteria at 1.5 h following phagocytosis by RAW264.7, differentiated THP-1, and iBMDM cells after 5 h of ampicillin treatment; *n* = 4. (D) Time-lapse images of *Salmonella* sensitive cells released from macrophages during antibiotic killing; scale bar, 2 μm; t = time (min). (E) Time-lapse images of persister cells with initial HslU-EGFP foci released from macrophages during antibiotic killing and subsequent resuscitation of surviving cells; scale bar, 2 μm; t = time (min). (F) Percentage of aggresome-containing bacterial cells in different subgroups of *Salmonella* released from macrophages (antibiotic-sensitive cells and persisters). (G) Contribution of aggresomes to persisters induced by macrophage phagocytosis. Orange denotes persister cells. The orange area in the red zone shows bacteria with protein aggresomes contributing to persister formation, while the orange in the blue zone represents those without aggresomes. (H) KEGG pathway analysis of macrophage phagocytosis-induced insoluble proteins isolated from *Salmonella*. (I) Live-cell imaging of *Salmonella* aggresomes and representative QUEEN 7μ_A81D 405ex and 488ex images of *Salmonella* in RAW264.7 cells at 1.5 h.p.i. (scale bar, 5 μm). (J) Average ATP levels in intracellular *Salmonella* with and without aggresomes. (K) Heat map displaying differentially expressed genes in *Salmonella*. (L) Heat map displaying differentially expressed genes between ROS-treated *Salmonella* and control samples; *n* = 3. (M) Fluorescence images of *sicA* promoter activity in intracellular *Salmonella* (scale bar, 5 μm). (N) Histogram showing *sicA* promoter activity in intracellular bacteria with and without aggresomes (O) Representative FACS plots of CellROX in infected macrophages at the indicated time points. (P) Representative FACS plots of mCherry signal from the P*sifA*_mScarlet-I reporter in *Salmonella* at 1.5 and 24 h.p.i. C was assessed using one-way ANOVA followed by Fisher’s LSD post hoc test; B, J and N were assessed using two-tailed unpaired *t*-test. Error bars indicate standard deviation; * *p* < 0.05, ** *p* < 0.01, and *** *p* < 0.001.

We then investigated the role of stress-induced aggresomes and found that ROS-induced aggresomes also promote persister formation (Figure S2A). Moreover, inhibiting aggresomes with macrophage ROS inhibitors restored antibiotic sensitivity in macrophage-released *Salmonella* (Figure S2B−D). *In vivo*, wild-type and *Cybb*^*-/-*^ mice were intragastrically inoculated with *S. Typhimurium* SL1344 and administered with 150 mg/kg cefotaxime intraperitoneally every 12 h. After 48 h, *Salmonella* persister ratio in the organs of *Cybb*^*-/-*^ mice was substantially lower than that in wild-type mice (Figure S2E).

Given the controversy regarding (p)ppGpp’s contribution to persister formation [9, 11], we reexamined it’s function in macrophage-induced *Salmonella* persistence. Using Δ*relA*Δ*spoT* ((p)ppGpp synthases), Δ*lon*, and Δ*lon*Δ*sulA* mutants of *S. Typhimurium* SL1344 [16], we found that these mutants showed no substantial decrease in the macrophage-induced persister cell ratio under ampicillin treatment (Figure S2F). BMDMs were isolated and differentiated by L929 supernatant, with flow cytometry analysis confirming over 90% macrophage purity (CD11b^+^ F4/80^+^) (Figure S2G). We then repeated the persister assay with various antibiotics including ampicillin, carbenicillin, meropenem, cefotaxime and nalidixic acid sodium. The results also showed a consistent conclusion (Figure S2F,H).

Interestingly, we found that the Δ*lon*, and Δ*lon*Δ*sulA* mutants contained more aggresomes than the wild-type strain during the 16-h stationary phase (Figure S2I,J). Meanwhile, the Δ*lon*, and Δ*lon*Δ*sulA* mutants from the stationary phase exhibited higher persister ratios than the wild-type *Salmonella* under various antibiotic treatments (Figure S2K). We also found that bacterial protein aggresomes was effectively suppressed by glucose supplementation, consequently diminishing persister cell formation (Figure S2L,M). In conclusion, our findings offer an alternative explanation that aggresome formation contributes greatly to *Salmonella* antibiotic persistence.

### Aggresomes induced by macrophage phagocytosis facilitate bacterial cell dormancy

To characterize *Salmonella* aggresome composition, we collected *Salmonella* released from macrophages sorted by Fluorescence-activated cell sorting (FACS) (Figure S3A), and isolated their insoluble proteins. Mass-spectrometry analysis identified 779 proteins (Table S1), with Kyoto Encyclopedia of Genes and Genomes (KEGG) pathway analysis revealing enrichment in ribosome, carbon metabolism, biosynthesis of amino acids and oxidative phosphorylation (Figure 2H). Primary sequestered components included ribosomal proteins (rpsA, rplQ, and rplC) impairing translation and amino acid biosynthesis enzymes (asnB, trpB, and gltB) compromising metabolic homeostasis [3, 17]. Comparative proteomics demonstrated consistent aggresome composition across infection, prolonged culture, and ROS stress conditions (Figure S3B,C). Following ROS stress challenge, bulk bacterial ATP levels decreased by nearly 10-fold (Figure S3D). Since bacteria possessing aggresomes encountered higher ROS levels (Figure 1K), we hypothesized they would have lower ATP concentrations. To investigate single-cell ATP dynamics, we employed the QUEEN 7μ_A81D biosensor [18], first validating its responsiveness in *Salmonella* through CCCP and 2DG-induced ATP depletion (Figure S3E). Subsequent *in vivo* infection assays revealed lower ATP levels in bacterial cells containing aggresomes (Figure 2I,J).

### Intracellular *Salmonella* containing aggresomes exhibit a dormant but SPI-1 T3SS highly expressing phenotype

To further investigate the interplay between *Salmonella* and host cells, we performed dual RNA sequencing of infected macrophages (Figure S4A and Table S2). Macrophage transcriptomics demonstrated the upregulated expression of host resistance factor-related genes associated with ROS production, nutritional immunity, autophagy, and inflammasome activation (Figure S4B), which can reduce intracellular *Salmonella* metabolism and restrict bacterial proliferation [19]. *Salmonella* transcriptome showed compensatory upregulation of genes involved in bacterial protein quality control and antioxidant defense system (Figure 2K). In addition, numerous metabolism-related genes involved in glycolysis, the TCA cycle, oxidative phosphorylation, and amino acid metabolism were suppressed (Figure 2K), suggesting that intracellular bacteria enter a dormant state.

Nevertheless, *Salmonella* can survive and replicate within macrophages according to the expression of type III secretion systems (T3SSs) encoded on *Salmonella* pathogenicity island 1 (SPI-1) and SPI-2 [20]. The expression of *Salmonella* SPI-1 genes increased following phagocytosis by macrophages for 1.5 h compared with that seen in extracellular bacteria (Figure 2K). Moreover, intracellular *Salmonella* with aggresomes exhibited higher SPI-1-related promoter activity than *Salmonella* without aggresomes (Figure S4C, 2M,N). RNA-seq results from *Salmonella* exposed to hydrogen peroxide (H2O2) stress indicated that ROS serves as a signal to induce bacterial T3SS1 effectors (Figure 2L, S4D−F, and Table S3). After exposure to H2O2 stress, *Salmonella* possessing aggresomes also exhibited higher fluorescence intensity of SPI-1 promoters than those lacking aggresomes (Figure S4G–I). Taken together, these findings suggest that *Salmonella* with aggresomes still maintain viability.

### Decreased ROS production by macrophages facilitates the expression of *Salmonella* SPI-2 effectors and regrowth

Next, we explored under what conditions bacteria with aggresomes can exit dormancy and resuscitate. We investigated the dynamic changes in ROS levels within macrophages at 1.5 h.p.i. and 24 h.p.i. The result of flow cytometry analysis revealed a decay in ROS stress experienced by intracellular *Salmonella* (Figure 2O). Concurrently, we observed an upregulation in the promoter activity of SPI-2 effectors, including *sifA* and *sspH2*, which facilitated the intracellular proliferation of *Salmonella* (Figure 2P, S4J). In addition, we investigated additional time points and plotted ROS levels against SPI-2 promoter activity. These results provided stronger correlative evidence regarding the link between reduced ROS production and increased SPI-2 effector expression (Figure S4K,L).

We supposed that the expression of SPI-2 genes determines the moment at which *Salmonella* possessing aggresomes begin to regrow in macrophages. To verify this assumption, the expression of the T3SS2 *sifA* promoter fused to mScarlet-I and bacteria resuscitation were monitored in real time. *Salmonella* were phagocytosed by macrophages at an MOI of 5 to ensure that there was only one bacterium per macrophage. When the expression of SPI-2 genes increased, *Salmonella* aggresomes disassembled, and the bacteria resumed growth and replication (Figure S4M). Taken together, these results revealed that a reduction in ROS stress within macrophages initiates the expression of SPI-2 genes, accompanied by *Salmonella* aggresome disassembly and later the bacteria regrowth and replication.

## CONCLUSION

In conclusion, our study demonstrates that *Salmonella*-containing aggresomes released from macrophages evade antibiotic killing by entering a metabolically dormant state, yet retain the ability to resuscitate through regulating virulence gene expression, ultimately enabling recurrent infections.

## AUTHOR CONTRIBUTIONS

Conceptualization: Xiao Chen, and Fan Bai; Methodology: Kefan Fang, Xiao Chen, Bo Li, Yingxing Li, Tian Tian, and Yifan Zhao; Investigation: Xiao Chen, Kefan Fang, Weixin Ke, Linqi Wang, Yuehua Ke, and Jing Geng; Visualization: Xiao Chen and Kefan Fang; Funding acquisition: Fan Bai and Mark C. Leake; Supervision: Fan Bai; Writing: Xiao Chen, Kefan Fang, Mark C. Leake, and Fan Bai.

## ACKNOWLEDGMENTS

This work was supported by grants from the National Science Fund for Distinguished Young Scholars (T2125002) and the New Cornerstone Science Foundation through the XPLORER PRIZE to F.B and the Engineering and Physical Sciences Research Council EPSRC (EP/W024063/1, EP/Y000501/1) to M.L. We thank Drs. Hideyuki Yaginuma (The University of Tokyo) and Yasushi Okada (RIKEN Center for Biosystems Dynamics Research) for sharing the QUEEN 7μ_A81D plasmid; Drs. Chunyan Shan, Liqin Fu (National Center for Protein Sciences Beijing, Peking University), and Jing Li (Imaging Core Facility, Technology Center for Protein Sciences, Tsinghua University) for their assistance with microscopy imaging; Drs. Bin Yu (Core Facility, Center of Biomedical Analysis, Tsinghua University) and Yinghua Guo (National Center for Protein Sciences Beijing, Peking University) for their technical support related to flow cytometry analysis; Drs. Dong Liu and Qi Zhang (National Center for Protein Sciences Beijing, Peking University) for their help with mass spectrometry; and Dr. Chenyang Geng (Peking University High-throughput Sequencing Center) for her assistance with RNA fragment analysis. We apologize for not being able to cite additional work owing to space limitations.

## DATA AVAILABILITY STATEMENT

All data supporting the findings of this study are included in the main text and the supplementary materials. Raw dual RNA sequencing and RNA sequencing data from this study have been deposited in NCBI under accession number PRJNA1013683 (https://www.ncbi.nlm.nih.gov/bioproject/PRJNA1013683). The transcriptome data analysis and image processing code utilized in this study have been deposited in the DRYAD repository and are publicly accessible at https://datadryad.org/stash/share/jxa9kd8pdEda_4SDRL2Z811D23I9TQoUr9CLoGErUHc. Supplementary materials (methods, figures, tables, graphical abstract, slides, videos, Chinese translated version and update materials) may be found in the online DOI.

## ETHICS STATEMENT

The ethics application was approved by the Research Ethics Committee of the Institute of Microbiology, Chinese Academy of Sciences, Beijing (No. SQIMCAS2020148).

## SUPPORTING INFORMATION

The online version contains supplementary methods, figures, and tables available.

Figure S1 Fluorescence labeling of bacteria and aggresomes.

Figure S2 ROS induces the formation of *Salmonella* aggresomes.

Figure S3 Correlation between *Salmonella* aggresome formation and bacterial antibiotic persistence.

Figure S4 Aggresomes induced by macrophage phagocytosis facilitate bacterial cell dormancy.

Figure S5 ROS is an activation signal for *Salmonella* SPI-1 genes.

Table S1 Total insoluble protein mass-spectrometric analysis.

Table S2 Dual RNA-seq data.

Table S3 Bacterial RNA-seq data.

Table S4 Strains, plasmids and primers used in this study.

## Materials and Methods

### Bacterial strains and plasmids

The *Salmonella Typhimurium* strains SL1344 and *Shigella flexneri* 2a strain 301 were grown in Luria Broth (LB) medium. *Salmonella* were inoculated into LB medium at a 1:1000 dilution and cultured for 16 hours to reach the stationary phase. Overnight cell cultures were transferred at a ratio of 1:100 into fresh LB medium and cultured for 3 hours to reach the exponential phase. *Mycobacterium smegmatis* strain mc^2^155 was grown in 7H9 broth medium. As required, antibiotics were added at the following concentrations: ampicillin 100 μg/mL, kanamycin 100 μg/mL, and chloramphenicol 25 μg/mL. Bacterial strains, plasmids, and DNA primers are described in Table S4.

### Construction of strains

Strains harboring target gene-fluorescent protein (FP) translational fusion or single-gene knockout mutants at the chromosomal locus were generated through gene replacement using the λ-red method. The kanamycin resistance gene was amplified from the pKD13 plasmid and introduced via electroporation into bacteria expressing the pSIM6 plasmid containing the λ-red recombinase [1, 2]. The kanamycin resistance gene flanked by frt sites was removed by expressing Flp recombinase using the pCP20 plasmid [1].

### Cell culture

RAW264.7 macrophages and THP-1 cells were purchased from the Cell Resource Center, Chinese Academy of Medical Sciences and Peking Union Medical College. The iBMDM cell line was a gift from the Zhengfan Jiang lab (School of Life Sciences, Peking University). RAW264.7 and iBMDM cells were cultured in Dulbecco’s modified Eagle’s medium (DMEM) supplemented with 10% fetal calf serum (FCS) at 37°C in a 5% CO_2_ incubator. THP-1 cells were cultured in Roswell Park Memorial Institute (RPMI) 1640 medium supplemented with 10% fetal calf serum (FCS) at 37°C in a 5% CO_2_ incubator. To differentiate monocytes into macrophage-like cells, THP-1 cells were cultured in 24-well plates at a density of 1 × 10^5^ cells per well in RPMI supplemented with 200 nM phorbol 12-myristate 13-acetate (PMA; Sigma) for 2 d. Bone marrow-derived macrophages were isolated from C57BL/6 mice and differentiated by supplementation with L929 supernatant.

### Bacterial infection of macrophages

For infection of mammalian cells, RAW264.7 macrophages were seeded onto 24-well plates (1 × 10^5^ cells/well) and cultured for 24 h at 37°C in a 5% CO_2_ incubator. The bacterial culture was added to the cells at an appropriate MOI and centrifuged at 800 × *g* for 5 min at room temperature (RT) to synchronize infection. After a 25-min incubation at 37°C, cells were washed three times with PBS to remove extracellular bacteria and incubated in fresh DMEM containing 100 μg/mL gentamycin for 1 h. Subsequently, cells were washed three times with PBS and incubated with fresh DMEM containing 50 μg/mL gentamicin for the remainder of the experiment.

### Mouse infection

WT C57BL/6 mice were purchased from Beijing Vital River Laboratory Animal Technology. B6.129S-Cybb^tm1Din^/J (*Cybb*^*-/-*^) mice were obtained from Jackson Laboratory [3]. All mouse experiments were carried out following the National Guidelines for Housing and Care of Laboratory Animals (Ministry of Health, China). The protocol conformed to institutional regulations after review and approval by the Institute of Microbiology, Chinese Academy of Sciences, Beijing. C57BL/6 mice (8 to12 weeks-old) were gavaged with 20 mg of streptomycin (in 200 mL sterile water), and 1 d later were inoculated intragastrically with 2 × 10^8^ *S. Typhimurium* strain SL1344 (in 200 μL of PBS). Bacterial inocula were prepared by 4 h of sub-culture in rich LB medium to reach mid-exponential growth phase prior to gavage. Water was offered immediately, and food was provided 2 h post-infection. Mice were euthanized at 2 h.p.i. and 120 h.p.i.

### FITC staining

*S. flexneri* and *M. smegmatis* were harvested and washed three times with PBS and incubated with 40 μg/mL FITC for 30 min [4].

### Cell imaging

After infection, macrophages were fixed in 4% paraformaldehyde (PFA) for 20 min, permeabilized with 0.1% Triton X-100 in PBS (pH 7.4), and then blocked with 1% bovine serum albumin (BSA). Cells were incubated overnight with anti-EEA1 (ab109110, abcam) or anti-LAMP1 antibodies (ab208943, abcam) at 4°C, and incubated the following day with Alexa Fluor 647-conjugated secondary antibodies. The actin cytoskeleton was stained with Alexa Fluor 647-conjugated phalloidin (magenta), and nuclei were stained with DAPI. Immunofluorescence imaging was performed using the DeltaVision OMX SR imaging system in conventional mode (GE Healthcare, USA), which was equipped with a 100 × oil immersion objective (NA 1.49) and EMCCD. Live-cell images were acquired under an inverted microscope (Zeiss Observer Z1, Germany).

### Tissue immunofluorescence imaging

Mice were transcardially perfused with 10 mL of ice-cold PBS buffer (containing 10 U/mL heparin), followed by 35 mL of 4% PFA in PBS. Perfusion-fixed PPs (following careful removal of intestinal contents), MLNs (following careful removal of fat tissue), and spleen were prepared and post-fixed in 4% PFA overnight at 4°C. The next day, the tissue was dehydrated with 30% sucrose in PBS, snap-frozen, cut into 8-μm sections using a cryostat, and placed on glass slides. Subsequently, slides were dried, fixed in 4% PFA for 20 min, blocked with 2% BSA, and permeabilized with 0.3% Triton X-100 in PBS (pH 7.4). Slides were incubated overnight with anti-F4/80 (14-4801-82, Invitrogen) at 4°C. The next day, slides were rinsed in PBS and incubated with Alexa Fluor 647-conjugated secondary antibodies (1:500) for 1 h (with 5-min staining with DAPI) at RT in the dark.

### Flow cytometry and cell sorting

Flow cytometry-based analysis was used to measure ROS levels in macrophages. Macrophages were seeded onto six-well plates, cultured for 24 h, and treated with 160 μM menadione or stimulated with the *S. Typhimurium* SL1344 strain. The culture medium was replaced with serum-free DMEM, and CellROX reagent was then added at a final concentration of 5 μM and incubated for 30 min at 37°C. Next, cells were treated with 5% trypsin and resuspended in PBS containing 1% FBS for flow cytometry analysis.

For bacterial cell sorting, Triton X-100 (0.1%) was added to each well and incubated for 20 min at 37°C to selectively lyse macrophages and release the mCherry-expressing *Salmonella*. Electronic noise was excluded using SSC and FSC, and then bacterial samples were sorted according to red fluorescence using a BD FACS Aria SORP Flow Cytometer. Subsequently, the sorted samples were prepared for the persister assay and mass spectrometry. For macrophage cell sorting, infected macrophages were sorted by gating mCherry-positive (mCherry^+^) cells and then collected in tubes containing 20% RNAlater solution diluted in PBS.

### Time-lapse recording of antibiotics killing and subsequent bacterial resuscitation

The *Salmonella* sorted by FACS were enriched by centrifugation at 15,000 × *g* for 15 min at 4°C. Bacteria were then exposed to 100 μg/mL ampicillin in 90% M9 + 10% LB medium for 4 h at 37°C, which was then replaced with fresh 90% M9 + 10% LB. An FCS2 flow cell chamber system equipped with a temperature controller (Bioptechs) was applied for single-cell observation.

### Insoluble protein isolation and mass spectrometry

*Salmonella* sorted by FACS after infection were harvested by centrifugation at 5000 × *g* for 15 min at 4°C. The method for insoluble protein isolation from *Salmonella* was modified from Tomoyasu [5]. Pellets were dissolved in 40 mL of buffer A (10 mM potassium phosphate buffer, pH 6.5, 1 mM EDTA, 20% (wt/vol) sucrose, and 1 mg/mL lysozyme) and incubated for 30 min on ice. Cell lysates were mixed with 360 mL of buffer B (10 mM potassium phosphate buffer, pH 6.5, and 1 mM EDTA) and sonication while cooling. The pellet fractions were resuspended in 400 mL of buffer C (buffer B with 2% NP 40) to dissolve membrane proteins. Finally, the aggregated proteins were isolated by centrifugation and re-suspended in 50 mL of buffer B by brief sonication. These isolated proteins were analyzed by mass spectrometry using an Orbitrap Fusion Lumos.

### Dual RNA-seq and data analysis

Infected macrophages were immediately fixed with 20% RNAlater solution in PBS before FACS. Total RNA was isolated from sorted infected cells using a mirVana kit (Thermo Fisher Scientific). The rRNA was removed using a Ribo-Zero plus rRNA depletion kit (Illumina) according to the manufacturer’s guidelines. cDNA libraries were generated using a VAHTS Universal V8 RNA-seq Library Prep Kit for Illumina (Vazyme). Sequencing was performed using the Illumina HiSeq platform.

Raw data were trimmed with Trimmomatic-0.39 to remove adapters, and the reads were then aligned to the *S. Typhimurium* SL1344 (NCBI RefSeq accession number: NC_016810.1) genome and the mouse (NCBI RefSeq accession number: GCF_000001635.26) genome using HISAT2 (2.2.1). Next, FeatureCounts 2.0.1 was used to generate a table of counts per gene in bacteria or macrophages for each sample for use in downstream differential expression analysis. Differential gene expression was analyzed using R 4.1.0. The “Control” for *Salmonella* infection refers to bacteria incubated in LB medium at 37°C to mid-exponential growth phase. The “Control” for macrophages refers to uninfected macrophages. The Z-score normalized values of genes were calculated for all control and dual RNA-seq samples and heatmaps were plotted using the pheatmap function in R 4.1.0.

### RNA-seq and data analysis

The bacterial cells were washed twice with normal saline and transferred to 2 mL of normal saline containing 25 mM H_2_O_2_ to induce ROS stress. Bacteria were harvested, and RNA was extracted for further analysis. For transcriptomics analysis, cDNA libraries were generated using the VAHTS Universal V8 RNA-seq Library Prep kit for Illumina (Vazyme). Sequencing was performed using an Illumina HiSeq platform. Sequencing adapter removal and quality-based trimming for raw data were performed using Trimmomatic v. 0.39 with TruSeq3 adapter sequences. Clean data were mapped to the reference genome using HISAT2 (2.2.1). Read numbers mapped to each gene were calculated using featureCounts. Finally, the count from each CDS was normalized to the FPKM value.

### ROS treatment

Exponential-phase and stationary-phase bacteria from LB cultures were washed twice with 0.9% NaCl and transferred to 2 mL of normal saline containing 25 mM H_2_O_2_ and 0.64 mM menadione, and incubated for 45 min at 37°C.

### Acid-shock treatment

Exponential-phase and stationary-phase bacteria from LB cultures were collected by centrifugation (4000 × *g* for 2 min at RT). Bacterial pellets were washed twice with 1 mL of PBS, resuspended in fresh acidified (pH 4.5) or neutral (pH 7.2) LB medium, and incubated for 30 min at 37°C.

### SHX treatment

Exponential-phase and stationary-phase bacteria from LB cultures were pre-exposed to 8 mM serine hydroxamate (SHX) in LB medium for 30 min.

### Persister assay

Bacteria released from macrophages were diluted 1:20 in fresh LB containing 100 μg/mL ampicillin and then incubated in a shaker (200 rpm) for 5 h at 37°C before estimating the colony forming unit (CFU) per milliliter. To evaluate the effect of macrophage-produced ROS on *Salmonella* persisters, macrophages were pretreated with 200 μM BHA, 50 μM VAS2870, 1 mM MnTBAP, 1 mM MitoTEMPO, and 5 μM mitoquinone for 3 h before performing the persister assay. There were four replicates.

Wild-type and *Cybb*^*-/-*^ C57BL/6 mice were intragastrically inoculated with 1 × 10^10^ CFU of *S. Typhimurium* SL1344 for acute systemic infection. After 2 days, the bacterial load in the spleen, liver, MLNs and PPs of the mice was evaluated. Infected mice were then administered with 150 mg/kg cefotaxime intraperitoneally every 12 h, and bacterial persisters were monitored after 48 h.

### Population-level measurement of bacterial ATP

Exponential-phase bacteria were washed twice with 0.9% NaCl, and then 25 mM H_2_O_2_ or 0.64 mM menadione was added for 45 min. Bulk ATP levels in the samples were measured using the BacTiter-Glo microbial cell viability assay. Intracellular ATP concentration was determined through the normalization of ATP levels by cell number and single-cell volume [6]. There were three replicates.

### Image analysis

Analysis of fluorescence imaging data was performed using Python 3.8.16. First, Omnipose 0.4.4 was applied to execute bacterial segmentation to obtain a mask for each bacterium [7]. Subsequently, the presence of protein aggresomes in the bacteria was determined based on the calculation of two fluorescence intensity parameters: maximum fluorescence intensity (fmax) and average fluorescence intensity (a). A bacterium was classified as a protein-aggregated cell if its maximum fluorescence intensity exceeded twice the average fluorescence intensity (fmax > 2a) and if there were a minimum of four adjacent pixels satisfying this criterion [8].

To determine the ATP sensor or ROS sensor ratio, the following steps were performed after characterizing protein aggresomes in each bacterium. First, the fluorescence channels at 405 nm and 488 nm were extracted from each bacterium, and the average fluorescence intensities of these channels were calculated separately. After subtracting the background fluorescence intensity, the ratio of the 405 nm / 488 nm fluorescence intensities in each bacterium was considered as the ROS sensor or ATP sensor ratio.

### Statistical analysis

Statistical analysis was performed using Origin Pro 2020b, as described in the figure legends. The mean is shown, with error bars indicating standard deviation (SD). The *p* value was calculated using two-tailed unpaired *t*-test or one-way ANOVA followed by Bonferroni/Fisher’s LSD post hoc test; \**p* < 0.05, \*\**p* < 0.01, \*\*\**p* < 0.001, and ns, not significant.

**FIGURE S1.**
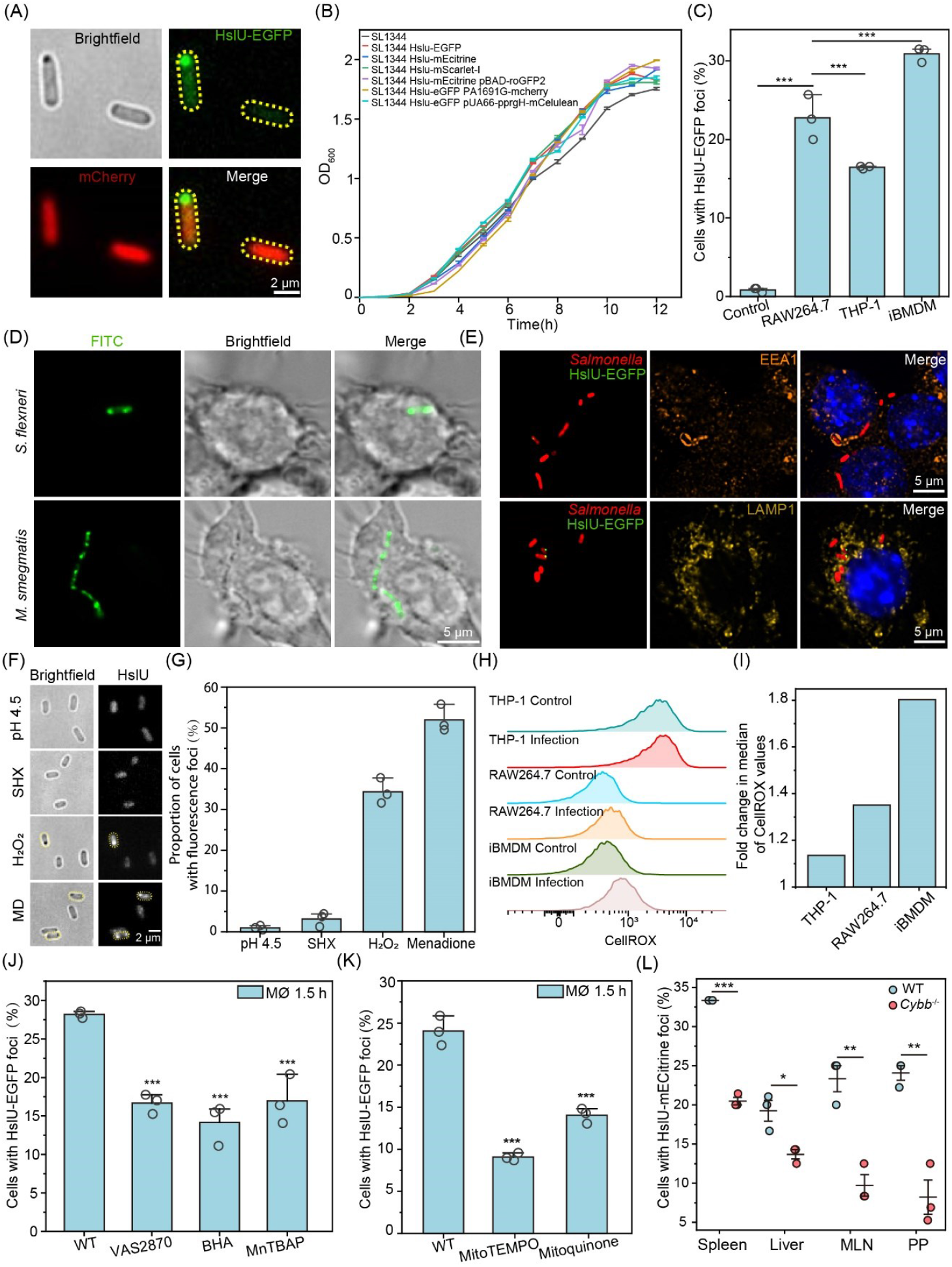
Fluorescence labeling of bacteria and aggresomes. (A) Brightfield and fluorescence images of cells labeled with a dual fluorescent protein reporter. *Salmonella* aggresomes were labeled with HslU-EGFP, and the bacterial cytoplasm was labeled with mCherry (scale bar, 2 μm). (B) Growth curves of the strains used in this study. (C) Percentage of *Salmonella* in the exponential phase possessing HslU-EGFP foci after internalization by RAW264.7 macrophages, THP-1 cells, and iBMDM cells; *n* = 3. (D) Live-cell imaging of aggresomes visualized by FITC in *S. flexneri* and *M. smegmatis*, followed by macrophage phagocytosis (scale bar, 5 μm). (E) Immunofluorescence microscopy imaging of *Salmonella* aggresomes in phagosomes. Early endosomes were stained with anti-EEA1 (orange), and late lysosomes were stained with anti-LAMP1 (yellow) (scale bar, 5 μm). (F) Brightfield and fluorescence images of HslU-EGFP/mScarlet-I labeled *Salmonella* cells following treatment with an acidic pH, Serine hydroxamate (SHX), H_2_O_2_, and menadione (scale bar, 2 μm). (G) Percentage of *Salmonella* cells possessing fluorescence foci following treatment with an acidic pH, SHX, H_2_O_2_, and menadione; *n* =3. (H) Representative histograms illustrating CellROX fluorescence. (I) Fold change in median CellROX fluorescence values in differentiated THP-1, RAW264.7, and iBMDM cells following *Salmonella* internalization. (J, K) Percentage of *Salmonella* possessing HslU-EGFP foci after internalization by macrophages in the presence of ROS inhibitors (J) and mitochondrial ROS inhibitors (K); *n* = 3. (L) Percentage of *Salmonella* cells with HslU-mECitrine foci in the spleen, liver, MLNs, and PPs of wild-type and *Cybb*^*−*/−^ mice at 48 h.p.i.; *n* = 3. C, J, and K were assessed using one-way ANOVA followed by Fisher’s LSD post hoc test; L was assessed using two-tailed unpaired *t*-test. Error bars indicate standard deviation; * *p* < 0.05, ** *p* < 0.01, and *** *p* < 0.001.

**FIGURE S2.**
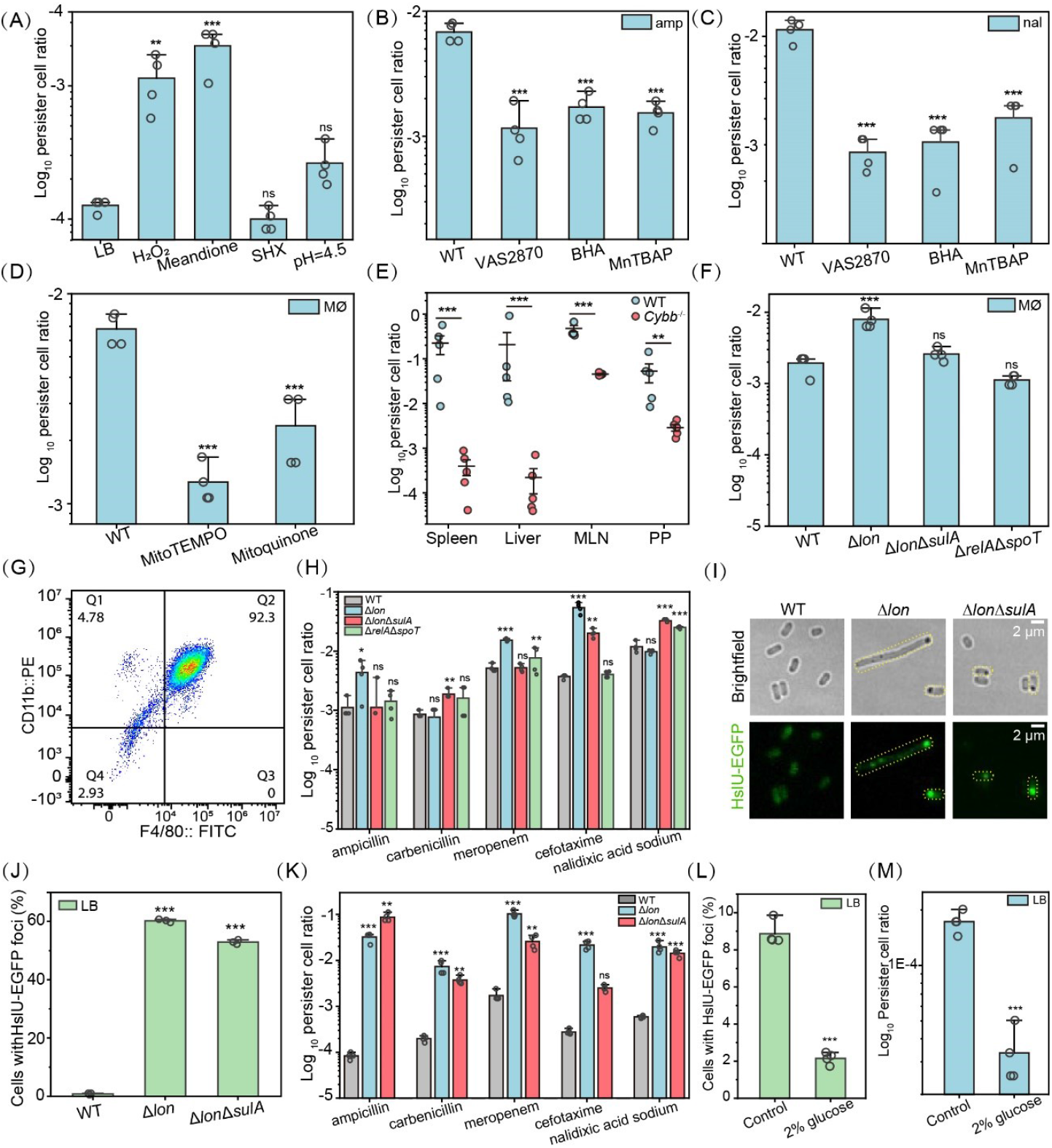
Correlation between *Salmonella* aggresome formation and bacterial antibiotic persistence. (A) Persister cell ratio of *Salmonella* SL1344 in the stationary phase subjected to an acidic pH, Serine hydroxamate (SHX), H_2_O_2_, and menadione stress; *n* = 4. (B) Persister cell ratio following treatment with 100 μg/mL ampicillin for *Salmonella* phagocytized by macrophages pretreated with ROS inhibitors; *n* = 4. (C) Persister cell ratio following treatment with 100 μg/mL nalidixic acid sodium for *Salmonella* phagocytized by macrophages pretreated with ROS inhibitors; *n* = 4. (D) Persister cell ratio following treatment with 100 μg/mL ampicillin for *Salmonella* phagocytized by macrophages pretreated with mROS inhibitors; *n* = 4. (E) *Salmonella* persister cell ratio recovered from the spleen, liver, MLNs, and PPs of wild-type and *Cybb*^*−*/−^ mice after 48 h of treatment with cefotaxime; *n* = 5. (F) Macrophage-induced persister cell ratio for the SL1344 wild-type and Δ*lon*, Δ*lon*Δ*sulA*, and Δ*relA*Δ*spoT* mutant strains 1.5 h after internalization by RAW264.7 macrophages; *n* = 4. (G) Flow cytometry analysis of BMDMs. Differentiated macrophages were stained with anti-F4/80 FITC and anti-CD11b PE. (H) Macrophage-induced persister cell ratio for the wild-type and Δ*lon*, Δ*lon*Δ*sulA*, and Δ*relA*Δ*spoT* mutant strains 1.5 h after internalization by BMDMs; *n* = 4. (I) Brightfield and fluorescence images of SL1344 wild-type and Δ*lon* and Δ*lon*Δ*sulA* mutant strains during the 16-h stationary phase (scale bar, 2 μm). (J) Percentage of *Salmonella* cells possessing HslU-EGFP foci in the SL1344 wild-type and Δ*lon* and Δ*lon*Δ*sulA* mutant strains; *n* = 3. (K) *Salmonella* persister cell ratio for the SL1344 wild-type and Δ*lon* and Δ*lon*Δ*sulA* mutant strains during the 16-h stationary phase; *n* = 4. (L) Aggresome formation in *Salmonella* SL1344 wild-type strains, with or without 0.2% glucose supplementation, as quantified by the percentage of bacterial cells containing HslU-EGFP foci; *n* = 4. (M) Persister cell frequency in SL1344 wild-type strains, with or without 0.2% glucose supplementation, following treatment with 100 μg/mL ampicillin; *n* = 4. A−D, F, H, J and K were assessed using one-way ANOVA followed by Fisher’s LSD post hoc test; E, L and M was assessed using two-tailed unpaired *t*-test. Error bars indicate standard deviation; * *p* < 0.05, ** *p* < 0.01, and *** *p* < 0.001.

**FIGURE S3.**
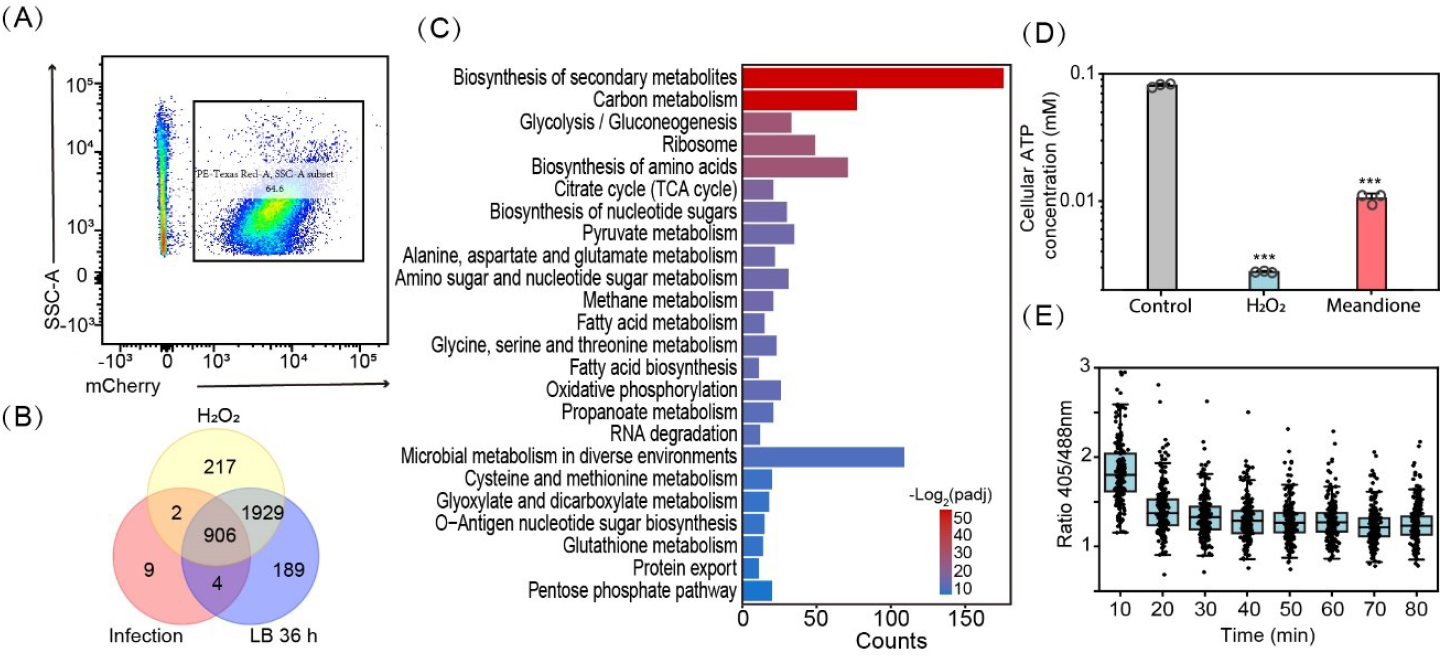
Aggresomes induced by macrophage phagocytosis facilitate bacterial cell dormancy. (A) Identification of *Salmonella* in macrophage-derived samples by flow cytometry. Bacterial cells were identified by first gating on events with similar FSC and SSC properties to *Salmonella*, followed by a second gating on particles emitting high levels of red fluorescence (bacterial cells constitutively expressing mCherry). (B, C) Venn diagram (B) and KEGG pathway analysis (C) showing the overlap of insoluble protein expression in bacteria cultured for 36 h in LB medium, subjected to H_2_O_2_ treatment and isolated from macrophage. (D) Cellular ATP concentration in cells treated with 25 mM H_2_O_2_ and 0.64 mM menadione for 45 min; *n* = 3. (E) Changes in the ratio of the QUEEN 7μ_A81D reporter (405 ex/488 ex) per single cell in stationary-phase cultures cotreated with 2-DG and CCCP. D was assessed using one-way ANOVA followed by Fisher’s LSD post hoc test. Error bars indicate standard deviation; *** *p* < 0.001.

**FIGURE S4.**
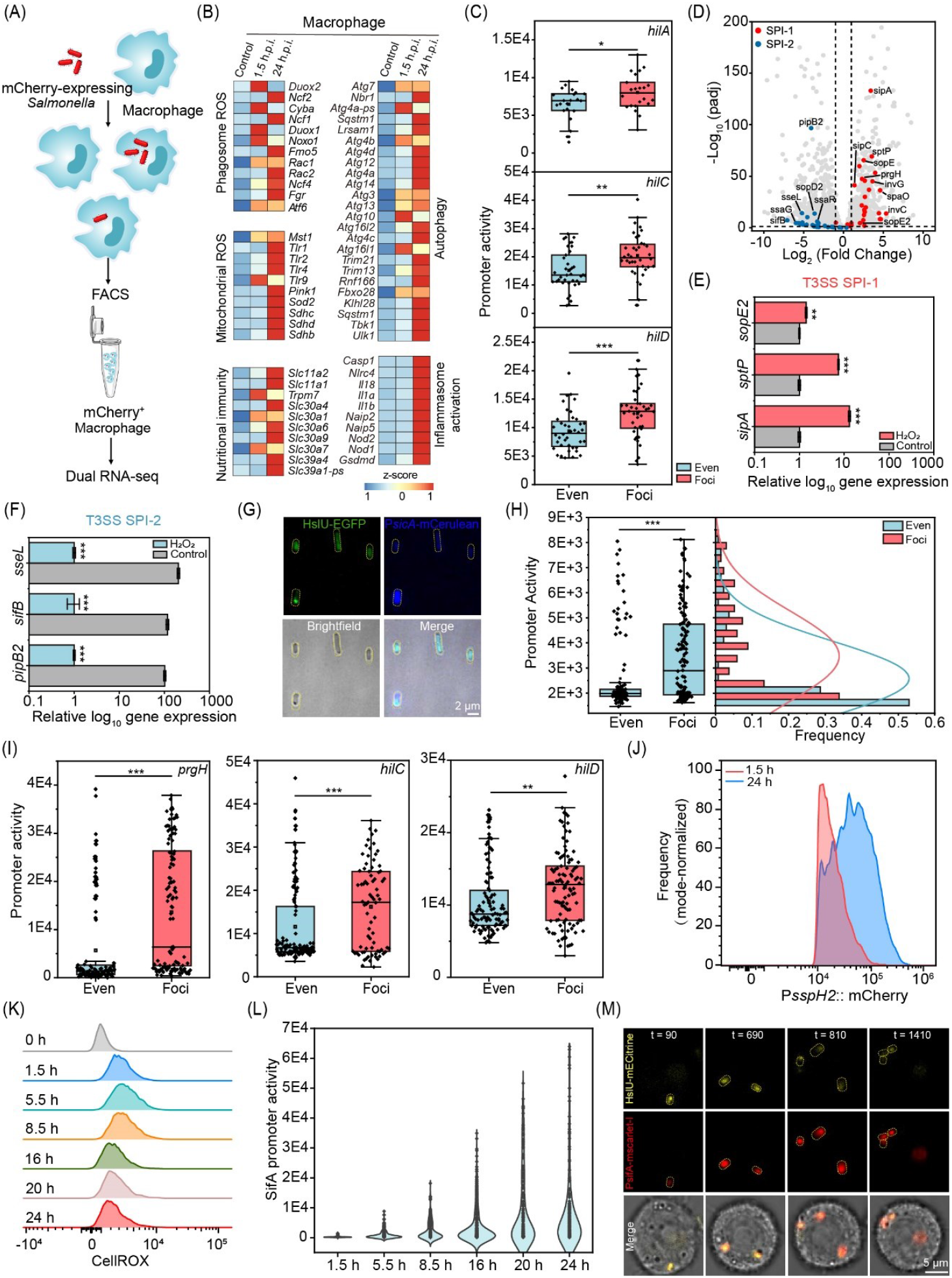
ROS is an activation signal for *Salmonella* SPI-1 genes. (A) Schematic summarizing the experimental design for dual RNA-seq. (B) Heat map displaying differentially expressed genes in macrophages. (C) SPI-1 T3SS promoter activity of SPI-1, including *hilA, hilC*, and *hilD* in intracellular *Salmonella* with and without aggresomes. (D) Volcano plots of differentially expressed mRNA; red dots, representative upregulated SPI-1 T3SS genes; blue dots, representative downregulated SPI-2 T3SS genes. (E, F) Fold increase in mRNA levels of SPI-1 T3SS (E) and SPI-2 T3SS (F) genes following exposure to 25 mM H_2_O_2_ compared with the saline control samples; *n* = 4. (G, H) Images (G) and histogram of *sicA* promoter activity (H) in bacteria following H_2_O_2_ treatment (scale bar, 2 μm). (I) SPI-1 T3SS promoter activity of *prgH, hilC*, and *hilD* at the single-cell level in *Salmonella* with and without aggresomes subjected to ROS-mediated stress. (J) Representative FACS plots of mCherry signal from the P*sspH2* mScarlet-I reporter in *Salmonella* at 1.5 and 24 h.p.i. (K) Representative FACS plots of CellROX in infected macrophages at the indicated time points. (L) Promoter activity of *sifA* in intracellular bacteria at the indicated time points. (M) Fluorescence images showing the replication of intracellular *Salmonella* with aggresomes and the promoter activity of *sifA*; scale bar, 5 μm; t = time (min). Two-tailed unpaired *t*-test was used for statistical analyses. Error bars indicate standard deviation; * *p* < 0.05, ** *p* < 0.01, and *** *p* < 0.001.

## REFERENCES

1. Balaban, Nathalie Q., Jack Merrin, Remy Chait, Lukasz Kowalik, Stanislas Leibler. 2004. “Bacterial persistence as a phenotypic switch.” Science 305: 1622−1625. 10.1126/science.1099390

2. Lewis, Kim. 2007. “Persister cells, dormancy and infectious disease.” Nature Reviews Microbiology 5: 48−56. 10.1038/nrmicro1557

3. Pu, Yingying, Yingxing Li, Xin Jin, Tian Tian, Qi Ma, Ziyi Zhao, Ssu-Yuan Lin, et al. 2019. “ATP-dependent dynamic protein aggregation regulates bacterial dormancy depth critical for antibiotic tolerance.” Molecular Cell 73: 143−156.e4. 10.1016/j.molcel.2018.10.022

4. Bollen, Celien, Elen Louwagie, Natalie Verstraeten, Jan Michiels, Philip Ruelens. 2023. “Environmental, mechanistic and evolutionary landscape of antibiotic persistence.” EMBO Reports 24: e57309. 10.15252/embr.202357309

5. Rowe, Sarah E., Nikki J. Wagner, Lupeng Li, Jenna E. Beam, Alec D. Wilkinson, Lauren C. Radlinski, Qing Zhang, Edward A. Miao, Brian P. Conlon. 2020. “Reactive oxygen species induce antibiotic tolerance during systemic Staphylococcus aureus infection. “ Nature Microbiology 5: 282−290. 10.1038/s41564-019-0627-y

6. Wilmaertsa, Dorien, Etthel M. Windels, Natalie Verstraeten, Jan Michiels. 2019. “General mechanisms leading to persister formation and awakening. “ Trends in Genetics 35: 401−411. 10.1016/j.tig.2019.03.007

7. Schumacher, Maria A., Kevin M. Piro, Weijun Xu, Sonja Hansen, Kim Lewis, Richard G. Brennan. 2009. “Molecular mechanisms of hipA-mediated multidrug tolerance and its neutralization by hipB.” Science 323: 396−401. 10.1126/science.1163806

8. Flannagan, Ronald S., Gabriela Cosío, Sergio Grinstein. 2009. “Antimicrobial mechanisms of phagocytes and bacterial evasion strategies. “ Nature Reviews Microbiology 7: 355−366. 10.1038/nrmicro2128

9. Helaine, Sophie, Angela M. Cheverton, Kathryn G. Watson, Laura M. Faure, Sophie A. Matthews, David W. Holden. 2014. “Internalization of Salmonella by macrophages induces formation of nonreplicating persisters.” Science 343: 204−208. 10.1126/science.1244705

10. Pontes, Mauricio H., Eduardo A. Groisman. 2019. “Slow growth determines nonheritable antibiotic resistance in Salmonella enterica. “ Science Signaling 12: eaax3938 10.1126/scisignal.aax3938

11. LeRoux, Michele, Peter H. Culviner, Yue J. Liu, Megan L. Littlehale, Michael T. Laub. 2020. “Stress can induce transcription of toxin-antitoxin systems without activating toxin.” Molecular Cell 79: 280−292.e8. 10.1016/j.molcel.2020.05.028

12. Jin, Xin, Ji-Eun Lee, Charley Schaefer, Xinwei Luo, Adam J. M. Wollman, Alex L. Payne-Dwyer, Tian Tian, et al. 2021. “Membraneless organelles formed by liquid-liquid phase separation increase bacterial fitness.” Science Advances 7: eabh2929. 10.1126/sciadv.abh2929

13. Li, Yingxing, Xiao Chen, Weili Zhang, Kefan Fang, Jingjing Tian, Fangyuan Li, Mingfei Han, et al. 2024. “The metabolic slowdown caused by the deletion of pspA accelerates protein aggregation during stationary phase facilitating antibiotic persistence. “ Antimicrobial Agents and Chemotherapy 68: e0093723. 10.1128/aac.00937-23

14. Ye, Yanfang, I-Ju Lee, Kurt W. Runge, Jian-Qiu Wu. 2012. “Roles of putative Rho-GEF Gef2 in division-site positioning and contractile-ring function in fission yeast cytokinesis.” Molecular Biology of the Cell 23: 1181−1195. 10.1091/mbc.E11-09-0800

15. Heijden, Joris van der, Else S. Bosman, Lisa A. Reynolds, B. Brett Finlay. 2015. “Direct measurement of oxidative and nitrosative stress dynamics in Salmonella inside macrophages.” Proceedings of the National Academy of Sciences of the United States of America 112: 560−565. 10.1073/pnas.1414569112

16. Harms, Alexander, Etienne Maisonneuve, Kenn Gerdes. 2016. “Mechanisms of bacterial persistence during stress and antibiotic exposure.” Science 354: aaf4268. 10.1126/science.aaf4268

17. Manhas, Reetika, Pankaj Tripathi, Sameena Khan, Bhavana Sethu Lakshmi, Shambhu Krishan Lal, Venkatraman Subramanian Gowri, Amit Sharma, Rentala Madhubala. 2014. “Identification and functional characterization of a novel bacterial type asparagine synthetase A: a tRNA synthetase paralog from Leishmania donovani. “ Journal of Biological Chemistry 289: 12096−12108. 10.1074/jbc.M114.554642

18. Yaginuma, Hideyuki, Shinnosuke Kawai, Kazuhito V. Tabata, Keisuke Tomiyama, Akira Kakizuka, Tamiki Komatsuzaki, Hiroyuki Noji, Hiromi Imamura. 2014. “Diversity in ATP concentrations in a single bacterial cell population revealed by quantitative single-cell imaging.” Scientific Reports 4: 6522. 10.1038/srep06522

19. Diacovich, Lautaro, Jean-Pierre Gorvel. 2010. “Bacterial manipulation of innate immunity to promote infection.” Nature Reviews Microbiology 8: 117−128. 10.1038/nrmicro2295

20. Haraga, Andrea, Maikke B. Ohlson, Samuel I. Miller. 2008. “Salmonellae interplay with host cells.” Nature Reviews Microbiology 6: 53−66. 10.1038/nrmicro1788

## Reference

1. Datsenko, Kirill A., Barry L. Wanner. 2000. “One-step inactivation of chromosomal genes in Escherichia coli K-12 using PCR products.” Proceedings of the National Academy of Sciences of the United States of America 97: 6640−6645. 10.1073/pnas.120163297

2. Datta, Simanti, Nina Costantino, Donald L. Court. 2006. “A set of recombineering plasmids for gram-negative bacteria.” Gene 379: 109−115. 10.1016/j.gene.2006.04.018

3. Matsumoto, Masanori, Seitaro Nakagawa, Lingzhi Zhang, Yuumi Nakamura, Amer E. Villaruz, Michael Otto, Christiane Wolz, Naohiro Inohara, Gabriel Núñez. 2021. “Interaction between Staphylococcus Agr virulence and neutrophils regulates pathogen expansion in the skin. “ Cell Host & Microbe 29: 930−940.e4. 10.1016/j.chom.2021.03.007

4. Li, Yanan, Weikang Liang, Chenghua Li. 2023. “Exogenous adenosine and/or guanosine enhances tetracycline sensitivity of persister cells.” Microbiological Research 270: 127321. 10.1016/j.micres.2023.127321

5. Tomoyasu, Toshifumi, Axel Mogk, Hanno Langen, Pierre Goloubinoff, Bernd Bukau. 2001. “Genetic dissection of the roles of chaperones and proteases in protein folding and degradation in the Escherichia coli cytosol.” Molecular Microbiology 40: 397−413. 10.1046/j.1365-2958.2001.02383.x

6. Conlon, Brian P., Sarah E. Rowe, Autumn Brown Gandt, Austin S. Nuxoll, Niles P. Donegan, Eliza A. Zalis, Geremy Clair, Joshua N. Adkins, Ambrose L. Cheung, Kim Lewis. 2016. “Persister formation in Staphylococcus aureus is associated with ATP depletion.” Nature Microbiology 18: 16051. 10.1038/nmicrobiol.2016.51

7. Cutler, Kevin J., Carsen Stringer, Teresa W. Lo, Luca Rappez, Nicholas Stroustrup, S. Brook Peterson, Paul A. Wiggins, Joseph D. Mougous. 2022. “Omnipose: a high-precision morphology-independent solution for bacterial cell segmentation.” Nature Methods 19: 1438−1448. 10.1038/s41592-022-01639-4

8. Pu, Yingying, Yingxing Li, Xin Jin, Tian Tian, Qi Ma, Ziyi Zhao, Ssu-Yuan Lin, et al. 2019. “ATP-dependent dynamic protein aggregation regulates bacterial dormancy depth critical for antibiotic tolerance. “ Molecular Cell 73: 143−156.e4. 10.1016/j.molcel.2018.10.022

